# Hybridization affects life-history traits and host specificity in *Diorhabda* spp

**DOI:** 10.1101/105494

**Authors:** E.V. Bitume, D. Bean, A.R. Stahlke, R.A. Hufbauer

## Abstract

Hybridization is an influential evolutionary process that has been viewed alternatively as an evolutionary dead-end or as an important creative evolutionary force. In colonizing species, such as introduced biological control agents, hybridization can negate the effects of bottlenecks and genetic drift through increasing genetic variation. Such changes could be beneficial to a biological control program by increasing the chances of establishment success. However, hybridization can also lead to the emergence of transgressive phenotypes that could alter host specificity; an important consideration when assessing potential non-target impacts of planned agents. In a series of lab experiments, we investigated the effects of hybridization between three species of *Diorhabda* released to control invasive *Tamarix* (saltcedar) on life history traits through two generations, and through the third generation for one cross. Depending on the cross, hybridization had either a positive or neutral impact on development time, adult mass, and fecundity. We evaluated preference for the target (saltcedar) relative to a non-target host *Tamarixaphylla* (athel), and found host specificity patterns varied in two of the three hybrids, demonstrating the possibility for hybridization to alter host preference. Importantly, the overall effects of hybridization were inconsistent by cross, leading to unpredictability in the outcome of using hybrids in biological control.

## 1. Introduction

Hybridization is an influential evolutionary process that has been viewed alternatively as an evolutionary dead-end, because hybrids are often less fit than the parental species (Mayr 1963; Dobzhanski 1970) or as an important creative evolutionary force (Anderson & Stebbins 1954; Ellstrand & Schierenbeck 2000). On the detrimental side, hybrid breakdown, or outbreeding depression, can decrease performance of hybrid individuals across a suite of traits linked to fitness, such as development time, mortality, and fecundity (Burton *et al.* 1999; Edmands 2002). On the positive side, hybridization can increase fitness relative to parents directly through heterozygote advantage (overdominance of beneficial traits) (Edmands 2002; Hedrick & Garcia-Dorado 2016; Lee *et al.* 2016) or by alleviating high mutational load (heterosis) and reducing inbreeding depression, and indirectly through restoring genetic variation lost through genetic drift or bottlenecks in population size (genetic rescue). Even in populations that have not experienced strong drift or bottlenecks, hybridization can increase overall population genetic variation, resulting in increased ability to respond to selection pressures (Fisher 1930). Additionally, hybridization can facilitate the formation of novel genotypes, potentially producing ‘transgressive’ phenotypes that fall outside the range of either parent (Rieseberg *et al.* 1999). Alternatively, hybridization can in some cases have minimal effects if the genetic distance between parents is small (Mallet 2005).

Intraspecific hybridization between recently diverged species may be particularly beneficial in colonizing populations that pass through strong bottlenecks in population size, in turn losing genetic variation, and potentially becoming inbred (Ellstrand & Schierenbeck 2000; Dlugosch & Parker 2008; Rius & Darling 2014; Laugier *et al.* 2016). In the planned release of specialized biological control agents, the goal is for the intentionally released population to establish and propagate (Seastedt 2015), to feed on the target host (typically an invasive weed or insect), and not shift to use other, non-target hosts. Biological control programs have a fairly low success rate (<50%), mostly due to lack of establishment of agents in their new environment (Van Driesche *et al.* 2010). As an evolutionary mechanism, hybridization might allow these establishing populations to better face adaptive challenges in the novel environment. There is some evidence that releasing different “strains” or ecotypes of biological control agents in an effort to increase genetic variation might improve establishment success (Hopper *et al.* 1993; Henry *et al.* 2010). New evidence suggests that increased genetic variation can be even more important than augmenting population size in promoting population growth (Frankham 2015; Hufbauer *et al.* 2015; Frankham 2016). Colonizing populations also experience novel environments in which transgressive phenotypes may, by chance, have higher fitness than parental phenotypes (Ellstrand & Schierenbeck 2000). Yet, the quantification of genetic variation in populations of biological control agents planned for release is not yet a standard procedure, likely because of the lack of studies investigating the effects of increased variation on long-term establishment.

Releasing genetically distinct ecotypes in the same area can promote hybridization. Only a few studies have looked at hybridization in biological control agents (Hoffmann *et al.* 2002; Mathenge *et al.* 2010; Benvenuto *et al.* 2012; Szucs *et al.* 2012). Szucs *et al.* (2012) found that hybridization improved performance in vital life-history traits, which could improve control of the target pest. However, there is also evidence that hybridization can decrease host specificity or increase host range as a result of changes in phenotype (Hoffmann *et al.* 2002; Mathenge *et al.* 2010). Such a change would increase the risk of biological control to non-target species dramatically. Thus, it is imperative that more experiments are executed to understand the consequences of hybridization for biological control programs, including evaluating the degree to which it will be possible to draw general conclusions versus research being needed on a case-by-case basis.

We present research in which we quantify the effects of hybridization between biological control agents in the genus *Diorhabda* that were released to control *Tamarix* (saltcedar, or tamarisk) in North America. Saltcedar in North America is comprised of a hybrid swarm of *Tamarix chinensis* and *T. ramosissima* (Gaskin and Schall 2002). It is an invasive weed that has colonized riparian habitats from Montana to Mexico (Gaskin & Schaal 2002). In 2001, the USDA’s Animal and Plant Health Inspection Service (USDA APHIS) approved the open field release of the central Asian salt cedar leaf beetle, *D. elongata*, as a biological control agent for saltcedar (DeLoach *et al.* 2003). *Diorhabda elongata* was classified as a dingle wide-ranging species that specialized on *Tamarix* spp and comprised different subspecies and ecotypes. To match environmental conditions in North America, the saltcedar biological control program eventually utilized seven *Diorhabda* ecotypes with native ranges stretching from north Africa to central Asia (Tracy & Robbins 2009; Bean *et al.* 2013a). A recent taxonomic revision of the *Tamarix*-feeding *Diorhabda* has used morphological and biogeographical data to define this group as a complex comprising five species: *D. meridionalis*, *D. carinulata*, *D. carinata*, and *D.sublineata* (Tracy & Robbins 2009). Recent genetic studies using amplified fragment length polymorphisms (AFLPs) revealed four major clades within this group which coincide with the four morphospecies (Bean *et al.* 2013b). There was also a fifth species, *D. meridionalis*, not currently used in the saltcedar biological control program. Currently, *D. carinulata* is the most widespread of the species in North America and covers large areas in Oregon, Idaho, Wyoming, Colorado, Utah, Nevada, northern Arizona, and northern New Mexico (Bean *et al.* 2013a). The other three species have all been released in Texas and have started spreading (Michels *et al.* 2013). Hybridization is possible between all four taxa, but hybrids between *D. carinulata* and each of the other three species produce few viable offspring. In contrast, egg viability of hybrids between *D. elongata*, *D. carinata*, and *D. sublineata* is comparable to that of the parents (Bean *et al.* 2013b). We crossed these three species reciprocally, and tracked performance over three generations to quantify the effects of hybridization. We measured several life history traits crucial to fitness, as well as host preference for saltcedar relative to a non-target plant, *Tamarix aphylla* (athel hereafter) for both hybrid offspring and the parental species.

## 2. Materials & Methods

### 2.1 Organism

The beetles used in our experiments stemmed from samples originally collected from saltcedar in Eurasia and North Africa. Descendants of these samples were used to establish laboratory populations maintained at the Palisade Insectary, Biological Pest Control Program, Colorado Department of Agriculture, Palisade, CO (CDA Palisade). Colonies were maintained on cuttings of saltcedar, including *T. ramosissima*, *T. chinensis*, and their hybrids (Gaskin & Schaal 2002), in 7.5 liter capacity plastic containers with mesh siding for ventilation in incubators under a light regime of 16:8 and 27°C/16°C. *Diorhabda carinata* (“C” hereafter) used in this study were originally collected in 2002 near Karshi (Qarshi), Uzbekistan (38.86 N, 65.72 E; elevation 350 m), and *Diorhabda sublineata* (“S” hereafter) originated near the town of Sfax, Tunisia (34.66 N, 10.67 E, elevation 10 m). Both these species were maintained in the lab prior to our experiments. *Diorhabda elongata* (“E” hereafter) were collected from Sfakaki, Crete, Greece (35.83 N, 24.6 E, elevation 7 m) and in 2004 they were first released upstream of Esparto, CA along Cache Creek in the Capay Valley. Unlike the other two species, *D. elongate* used in our experiments were collected in 2015 from the field in the Capay Valley and used to start a laboratory colony. No other species were released into the Capay Valley nor have any been established within 150 miles, and therefore it is reasonable to assume that there was no chance for hybridization before our experiments.

### 2.2 Crosses

To produce the first generation of hybrids, seven virgin females and seven males of each species were placed together into a plastic bucket with mesh siding (7.5 liter) with saltcedar. Since male female directionality can affect the fitness of hybrid offspring (Payseur & Rieseberg 2016), we crossed each species reciprocally. We thus made the following hybrids: *D. carinulata D. elongata* (C_f_ ×; E_m_, E×; C_m_), *D. carinulata* x *D. sublineata* (C_f_×; S_m_, S_f_ ×; C_m_), *D. elongata* x *sublineata* (E_f_ ×; S_m_, S_f_ ×; E_m_), plus the parents (C_f_ ×; C_m_, E_f_ ×; E_m_, S_f_ ×; S_m_,). To keep inbreeding depression to a minimum, we initiated two separate buckets for each of the parental lines so that density remained the same but so the parental generation had 14 families rather than 7 for the crosses. All adults were allowed to remain in the buckets for five days of egg-laying.

### 2.3 F_1_ adult performance test

We counted the number of eggs produced over 48 hours as an estimate of performance of first generation hybrids. Buckets were checked daily for emergence of *F_1_* adults. On the day of emergence, adults were sexed and mating pairs were placed into a plastic container (0.4L) with a paper towel lining the bottom and food. The containers were checked daily for eggs. The number of eggs produced was counted for 48 hours after the first eggs were laid. After this time, *F_1_* adults were removed and killed by freezing.

### 2.4 F_2_ larval performance test

We measured percent hatching of all eggs laid in the first 48 hours, development time (in days), and adult mass (mg) attained by each *F_2_* larva. Upon emergence, the date was recorded as well as the number of eggs that successfully hatched. Counting eggs is challenging due to the three-dimensional nature of the egg clutches. Following (Bean *et al.* 2013b), to ensure accuracy we also counted the number of larvae and compared this with the number of eggs. If the number of eggs was less than the number of larvae, we used the number of larvae as the total number of eggs produced. If the number of eggs was greater than the number of larvae, we conducted a recount of the clutch. Out of the hatched larvae from each mating pair, up to five were randomly chosen and allowed to develop individually.

Larvae were maintained in small plastic cups (0.4L) and given fresh saltcedar with its stem in a water-filled 1.5mL eppendorf tube each day. A paper-towel lined the bottom of each cup. When the larvae reached their last stage of development, 2 cm of sand was placed in each cup to provide conditions favorable for pupation. All larvae were maintained in incubators under a light regime of 16:8 (L: D) and 27°C/16°C, and rotated every other day to standardize environmental effects.

### 2.5 F_2_ adult preference test

We conducted a host preference test to determine if hybridization affected host preference for the non-target species, athel, presenting beetles with a choice between saltcedar and athel. Athel is an ornamental that is found at more southern latitudes in the US and is considered invasive in the southwestern U. S. (Gaskin & Shafroth 2005). *Tamarix* hybrids of *T.ramosissima* and *T. chinensis* (saltcedar) and are considered the preferred field host of *Diorhabda*. Previous host testing showed that the three *Diorhabda* species used in this study can survive as well on athel as on saltcedar, will oviposit on either saltcedar or athel under laboratory no-choice conditions, and showed an inconsistent preference for saltcedar under choice conditions (Milbrath & Deloach 2006a; Milbrath & DeLoach 2006b). However, saltcedar is preferred at field sites (Moran *et al.* 2009), and the intrinsic rate of increase of beetle populations is reduced on athel due to smaller egg mass size and a delayed start to oviposition (Milbrath & DeLoach 2006b).

Between 24 and 48 hours after emergence, we sexed and weighed the *F_2_* adults. The beetles were placed in a plastic tub (3L) with two eppendorf tubes containing equal amounts of either athel or saltcedar. Each beetle was placed in the middle of the tub, with both plants placed equidistantly at 10 cm from the center. The beetle remained in the plastic tub for 24 hours, at which time the amount of frass under each plant was weighed to the nearest 0.1 g (DeLoach *et al.* 2003).

### 2.6 F_3_ larval performance test on two different hosts

We measured *F_3_* larval performance on athel and saltcedar. After the host-choice test, mating pairs were formed with *F_2_* adults from the same cross. All *F_2_* adults were given saltcedar foliage to feed on regardless of what they chose as their host in the adult preference test. They were placed in the same plastic dish as previously described and allowed to mate and oviposit. The date of first oviposition, the number of eggs laid in 48 hours, and the percent hatching was recorded. Larvae from each mating pair were split and a maximum of five larvae were placed in a plastic dish with either athel or saltcedar. We measured development time to adult and adult mass.

### 2.7 Statistical analysis

Our interests center on comparing the fitness of hybrids to their parental species. Thus, each analysis was done separately for each of the 7 pairs of parental species and their respective two hybrid crosses (male/female reciprocal). All statistical analysis was conducted using R version 3.3.2 (R_Core_Team 2016). For the first generation, we analyzed differences in the total number of eggs produced between hybrids and parental species using a standard linear model. The number of eggs was log-transformed to meet the assumption of homogeneity of variance. Percent hatching was analyzed as a proportion of hatched eggs compared to the total number of eggs using a generalized linear model with quasibionmial error distribution. Cross was the only fixed effect for number of eggs produced and percent hatching in the first-generation analysis. For the second and third generations, we quantified the development time from egg to adult (days), adult mass (mg), and host choice. For development time and adult weight, we used linear mixed-effects models through the lme4 package (Bates *et al.* 2015) with cross, sex and their interaction as fixed effects, and family as a random effect. For host choice with a binary response (saltcedar or athel), we used a generalized linear mixed effects model with a binomial error distribution. For the third generation, we also included the random effect of cup nested within family for development time and adult weight.

## 3. Results

### 3.1 Egg count, percent hatching

Hybridization did not significantly affect the number of *F_1_* eggs laid in 48 hours for any of the crosses (Tables 1–3). Cross had a marginally significant influence on percent of eggs that hatched with the E×E and E_f_×Sm cross producing slightly fewer viable eggs than the other crosses (F_3_,37 = 2.82, *P* = 0.052, Table 2). In the *F_2_*, only for the S×C cross was there a significant effect of cross on the number of eggs laid in 48 hours, where hybrids produced significantly more eggs than either parental species (F_3_, 39 = 2.97, *P* = 0.044, Table 1, Figure 1).

**Figure 1:**
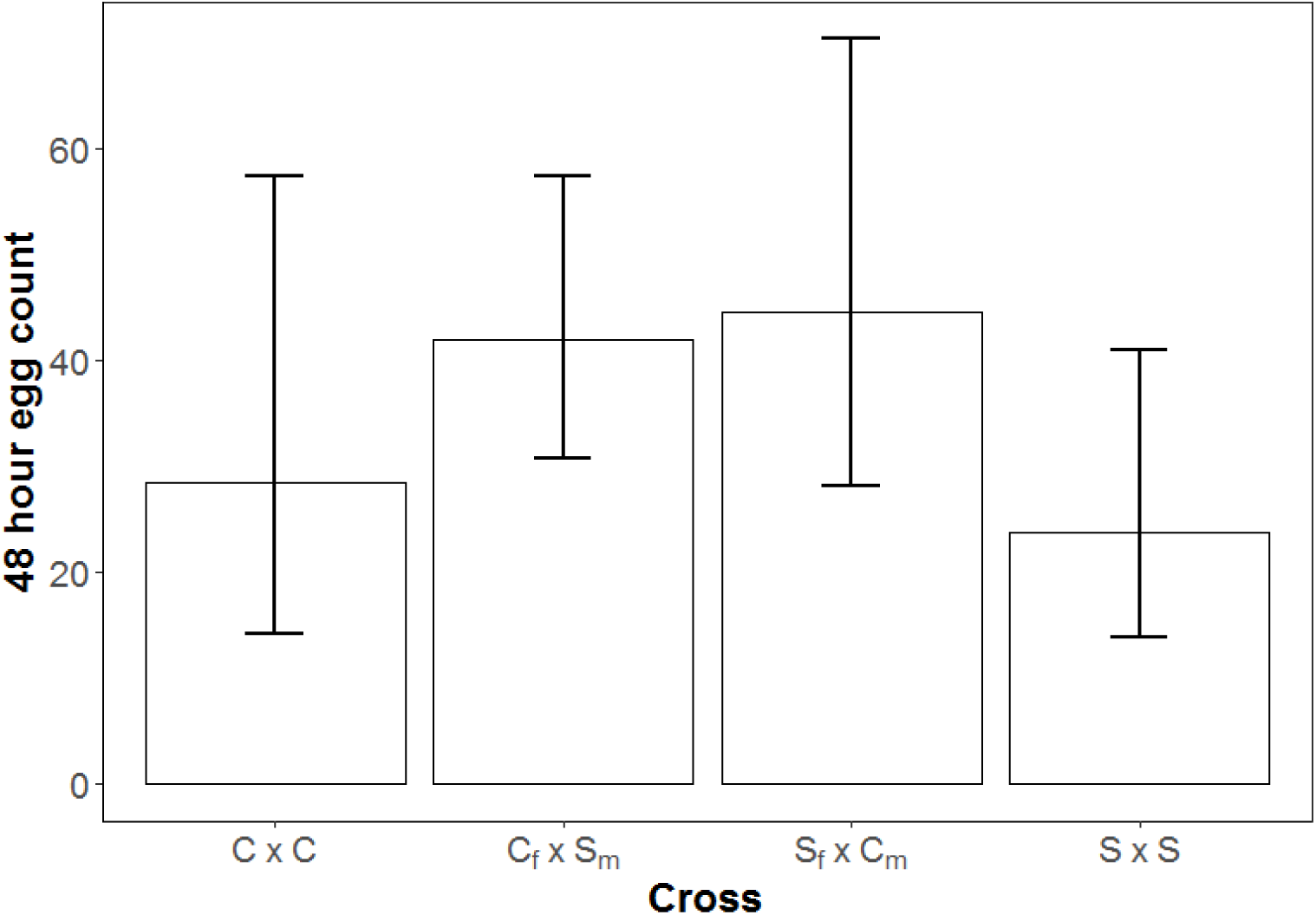
Number of eggs produced in first 48 hours by *D.carinata* (C×C), *D. sublineata* (S×S), and their hybrids in thesecond generation. Cross significantly affected egg production, with hybrids producing more than either parental species. Bars represent 95% confidence intervals

**Table 1:**
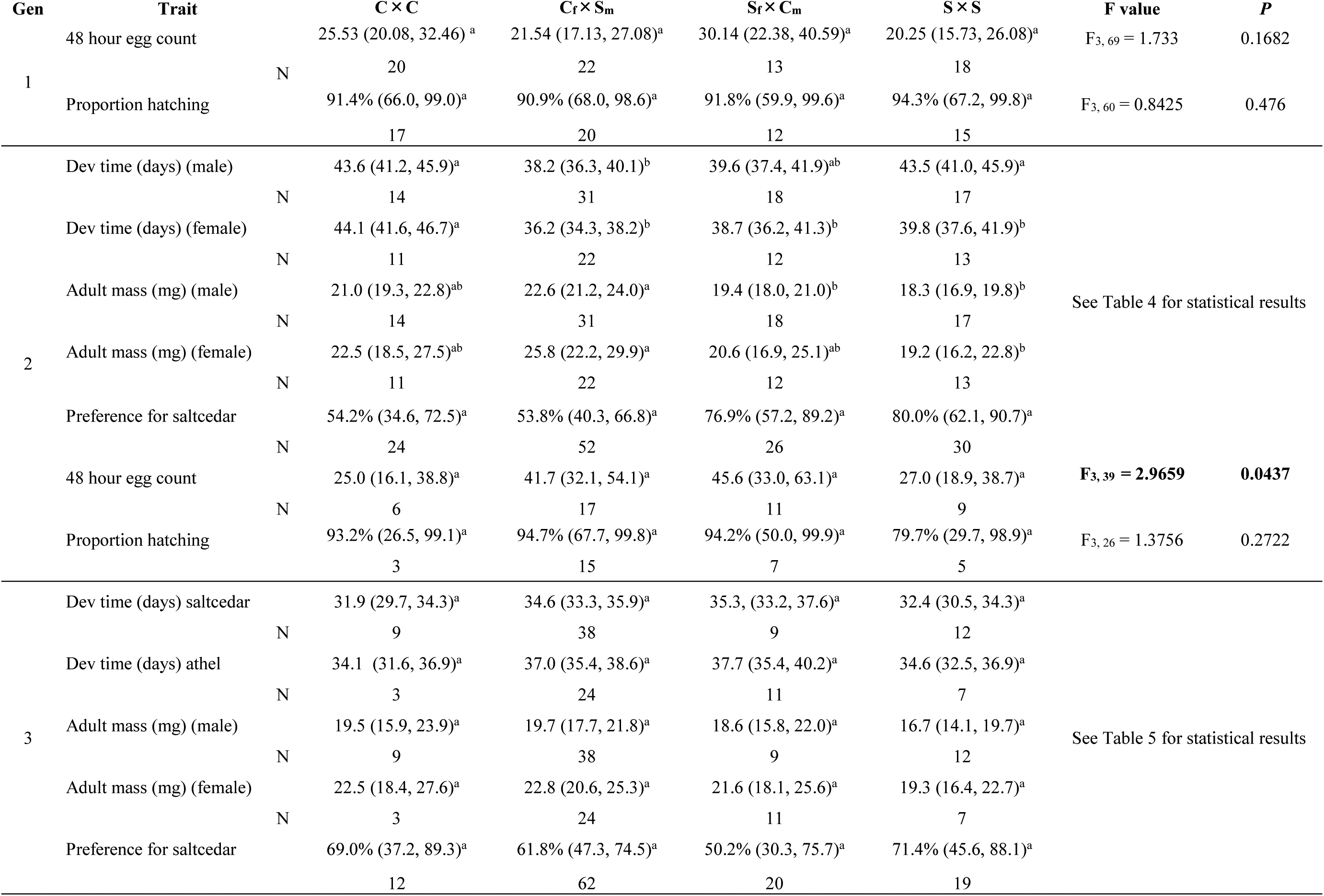
Trait means (95% CI) for each generation of the *D. sublineata* by *D. carinata* cross. Letters indicate significant differences between crosses.

**Table 2:**
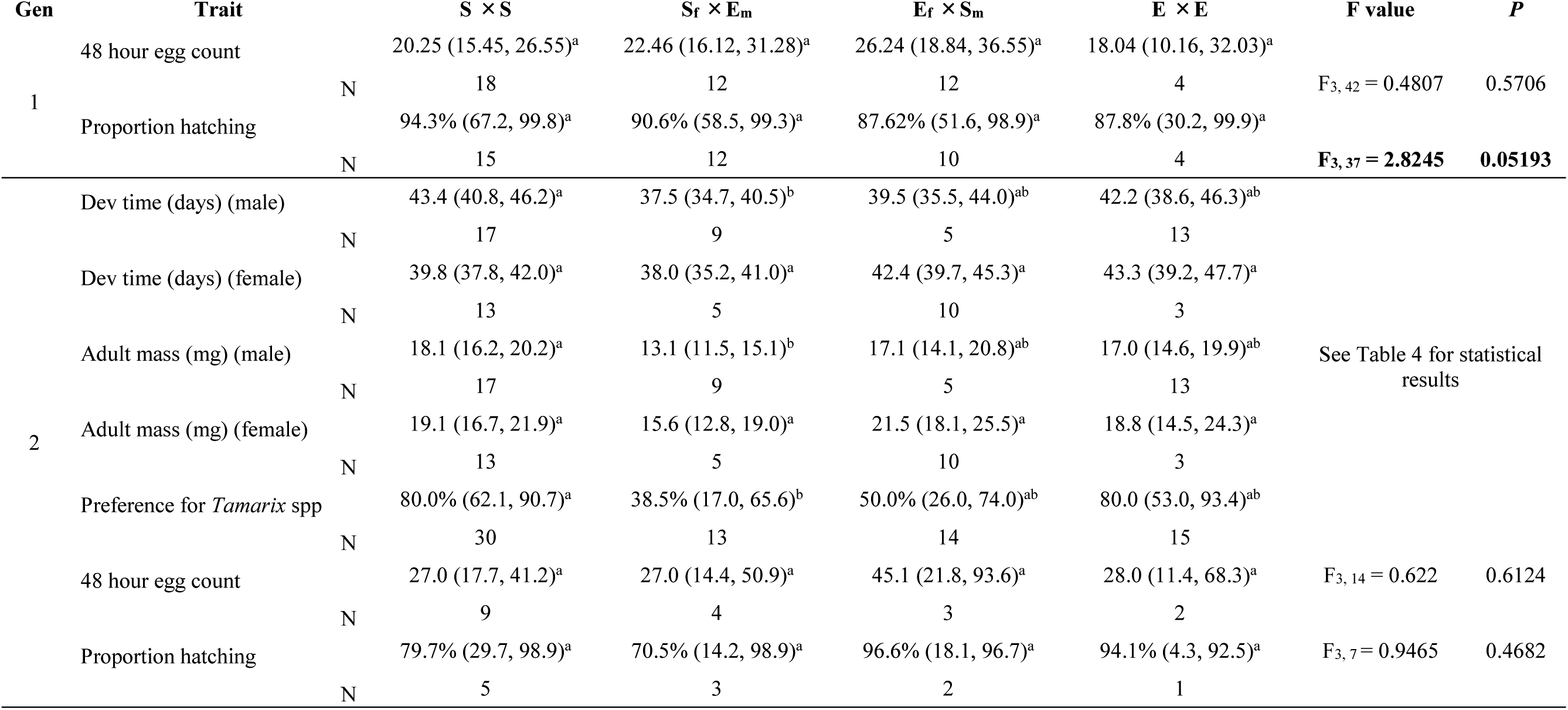
Trait means (95% CI) for each generation of the *D. sublineata* by *D. elongata* cross. Letters indicate significant differences between crosses. Cross

**Table 3:**
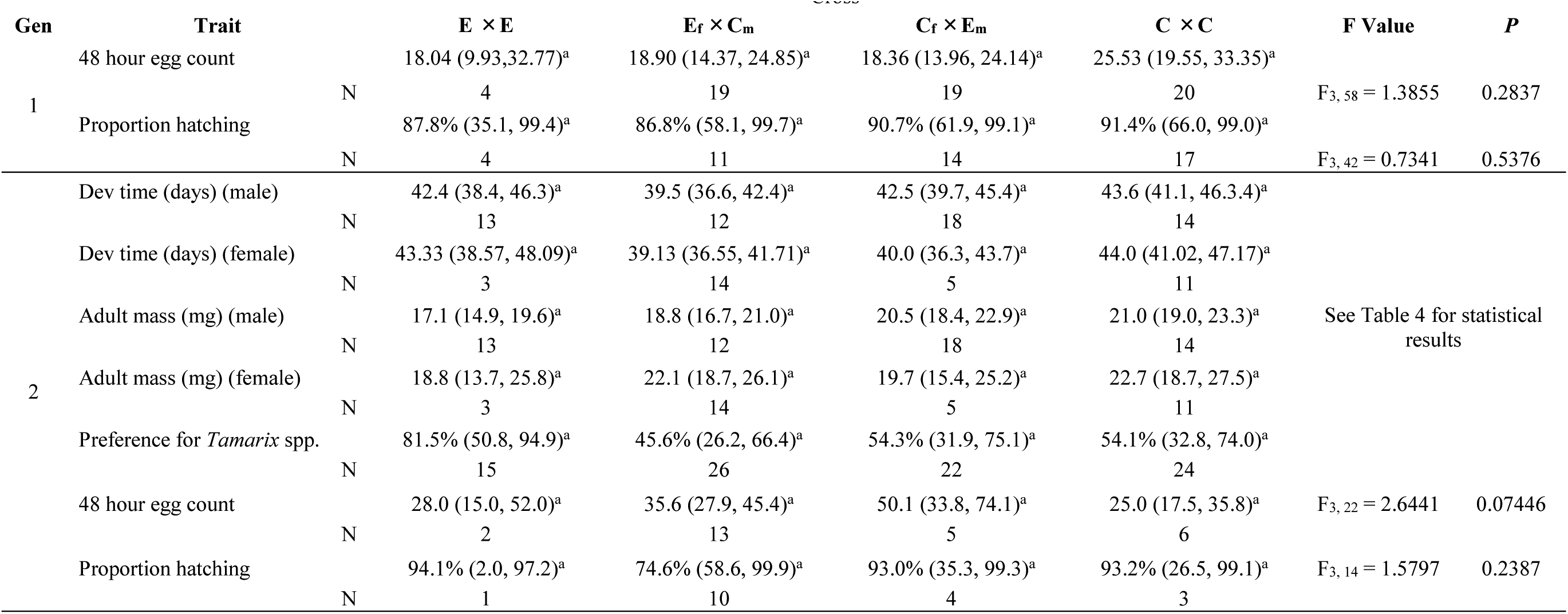
Trait means (95% CI) for each generation of the *D. elongata* by *D. carinata* cross. Letters indicate significant differences between crosses.

Cross did not affect the percentage of eggs hatched for any crosses in the second generation (Table 1–3).

### 3.2 Development time, adult mass

In the S×; C crosses, females were larger (effect of sex 
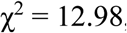
 df = 1, *P* <0.001, effect of cross: 
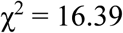
 df = 3, *P* <0.001) and hybrids developed faster (effect of sex: 
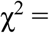
 9.93, df = 1, *P* = 0.002, effect of cross:
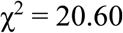
 df = 3, *P*<0.001) (Table 1, 4, Figure 2). For the S×E cross, there was a significant interaction between sex and cross in development time, in that males developed slower than females in the S×S line (interaction cross*sex: 
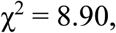
 df = 3, *P* = 0.031, Tables 2 and 4). We also found a significant effect of sex and cross on adult mass in the S×E cross, with females being overall larger (effect of sex: 
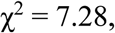
 df = 1, *P* = 0.007, effect of cross 
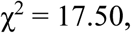
 df = 3, *P* <0.001, Tables 2 and 4). There was no effect of sex or cross on development time or adult mass for the E×C cross, although overall, females tended to be larger.

**Figure 2:**
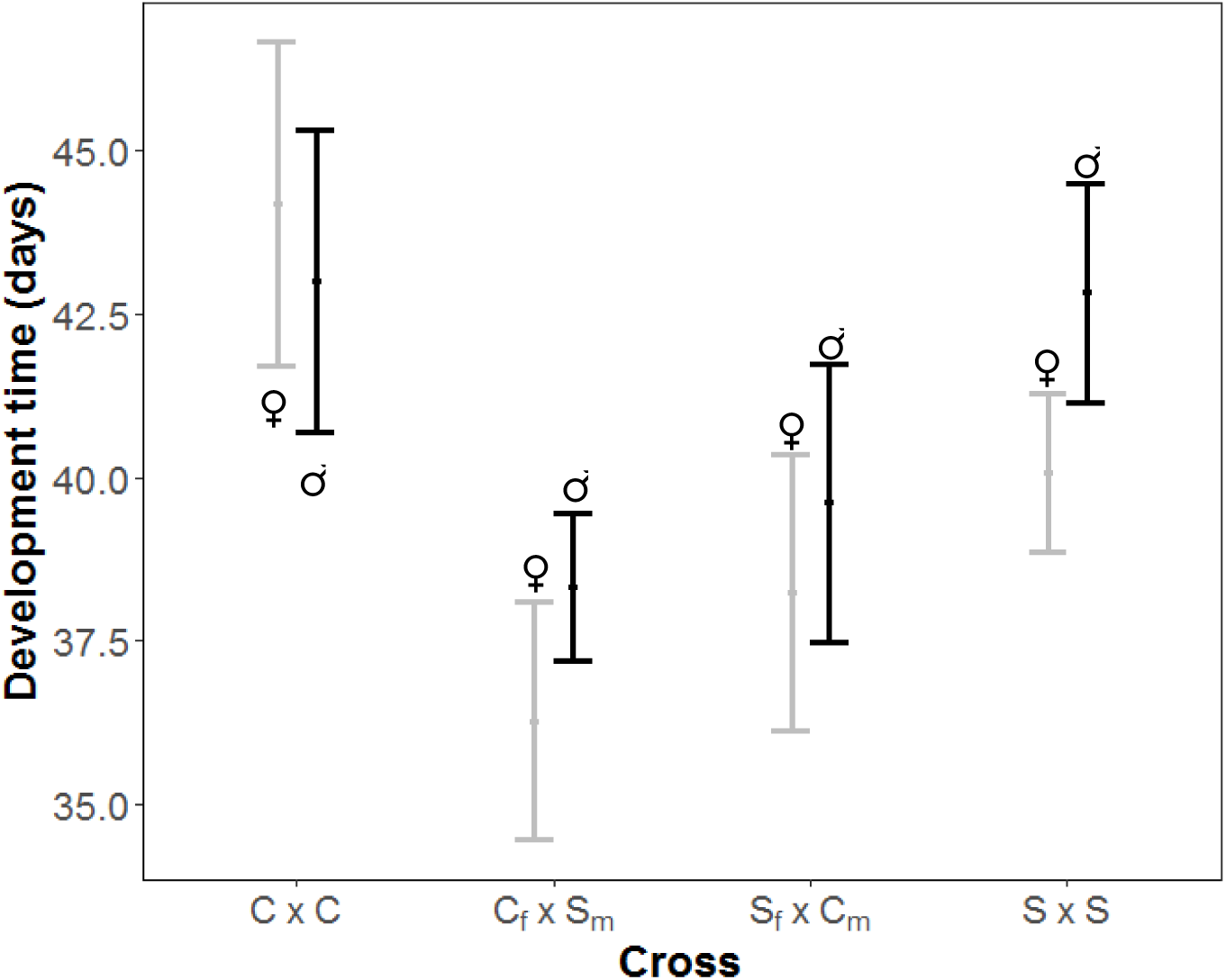
Development time from hatching until adult for each sex of *D. carinata* (C × C), *D. sublineata* (S × S) and their hybrids in the second generation. Cross significantly affected development time, with hybrids developing faster than either parental species. Grey and black lines represent females and males, respectively. Bars represent 95% confidence intervals.

**Table 4:** Results from generalized linear mixed-effects models for the second generation of hybridization for all crosses.

| Trait | **S × E Cross** |  |  | **Random effects** |  |  |  |
| --- | --- | --- | --- | --- | --- | --- | --- |
|  | Cross | Sex | Cross*sex | Family |  | Residual |  |
| | $\chi^2$, (df), *P* | $\chi^2$, (df), *P* | $\chi^2$, (df), *P* | Variance | Std dev | Variance | Std dev |
| Dev time | **10.82, (3), 0.013** | 0.47, (1), 0.493 | **8.90, (3), 0.031** | 0.006174 | 0.07857 | 0.001771 | 0.04208 |
| Adult mass | **17.50, (3), <0.001** | **7.28, (1), 0.007** | 2.56, (3), 0.464 | 0.01266 | 0.1125 | 0.02347 | 0.1532 |
| Preference for saltcedar | **9.23, (3), 0.026** | 0.98, (1), 0.322 | 0.99, (3), 0.804 | 0 | 0 | 0 | 0 |
| **C × E Cross** |  |  |  |  |  |  |  |
| Dev time | 5.99, (3), 0.112 | 0.08 (1), 0.7837 | 5.79, (3), 0.122 | 0.008246 | 0.09081 | 0.001845 | 0.04295 |
| Adult mass | 5.68, (3), 0.128 | 3.47, (1), 0.0626 | 2.65, (3), 0.449 | 0.01058 | 0.1029 | 0.0348 | 0.1868 |
| Preference for saltcedar | 3.66, (3), 0.300 | 0.89, (1), 0.3464 | 3.69, (3), 0.297 | 0.6548 | 0.8092 | 0 | 0 |
| **S × C Cross** |  |  |  |  |  |  |  |
| Dev time | **20.60, (3), <0.001** | **9.934, (1) 0.002** | 5.82, (3), 0.120 | 9.787 | 3.128 | 4.205 | 2.051 |
| Adult mass | **16.39, (3), <0.001** | **12.982, (1), <0.001** | 2.73, (3), 0.435 | 0.02041 | 0.1429 | 0.02266 | 0.1505 |
| Preference for saltcedar | **9.87, (3), 0.031** | 0.3953, (1), 0.5295 | 1.89, (3), 0.596 | 0 | 0 | 0 | 0 |

For the third generation, we were only able to investigate the effects of hybridization for the S × C cross (*D. sublineata* x *D. carinata*) due to limitations in the availability of our host plants. Development time was significantly affected by host plant and by cross, with both parents and hybrids developing slower on athel (effect of cross 
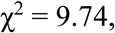
 df = 4, *P* = 0.029; effect of plant:
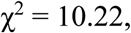
 df = 1, *P* = 0.001, Tables 1, 5, Figure 3). While there is a trend for hybrids to develop slower than parents regardless of host plant, there was no significant decrease in development time in hybrids compared to parents (effect of cross: Hybrids develop slower than the parents regardless of host plant, yet contrasts between crosses were not significant. Adult weight was not affected by hybridization, however females were larger regardless of cross (effect of sex 
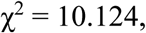
 df = 1, *P* = 0.001, Tables 1, 5).

**Figure 3:**
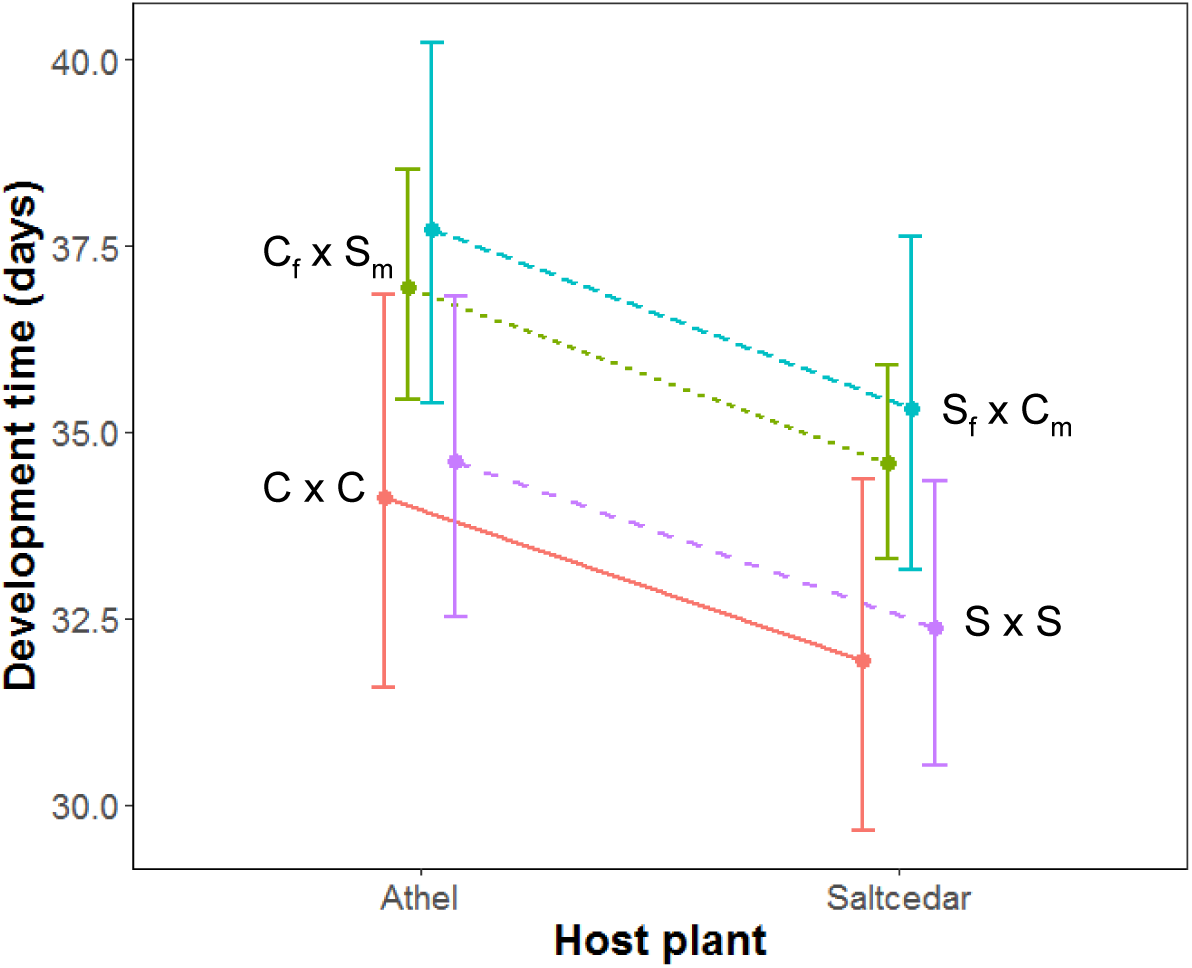
Development time on athel (non-target) and saltcedar (target) for *D. carinata* (C × C), *D. sublineata* (S × S), and their hybrids after three generations of hybridization. S_f_ × C_m_ are long close dashes, C_f_ × S_m_ small dashes, S × S long spaced dashes, and C × C solid line. Host plant significantly affected development time with beetles developing slower on the non-target host.

### 3.3 Host choice

We tested the host preference of individuals from all crosses in the second generation. Due to limitations in our host plant resources and, because of differences seen in the second generation, we also examined host preference for the S × C cross in the third generation. Sex did not affect host choice for any of the crosses in the second generation (Tables 1–3). Cross significantly affected host choice in the S × C cross (effect of cross: χ^2^=9.87 df = 3, *P* = 0.031) and S × E cross (effect of cross (χ^2^=9.23 df = 3, *P* = 0.026)whereas there was no difference in host preference between hybrids and their parents in the E × C cross (Tables 1–4, Figure 4). There was no effect of hybridization on host preference in the S × C cross in the third generation (effect of cross 
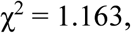
 df = 3, *P* = 0.7619, Tables 1, 5).

**Figure 4:**
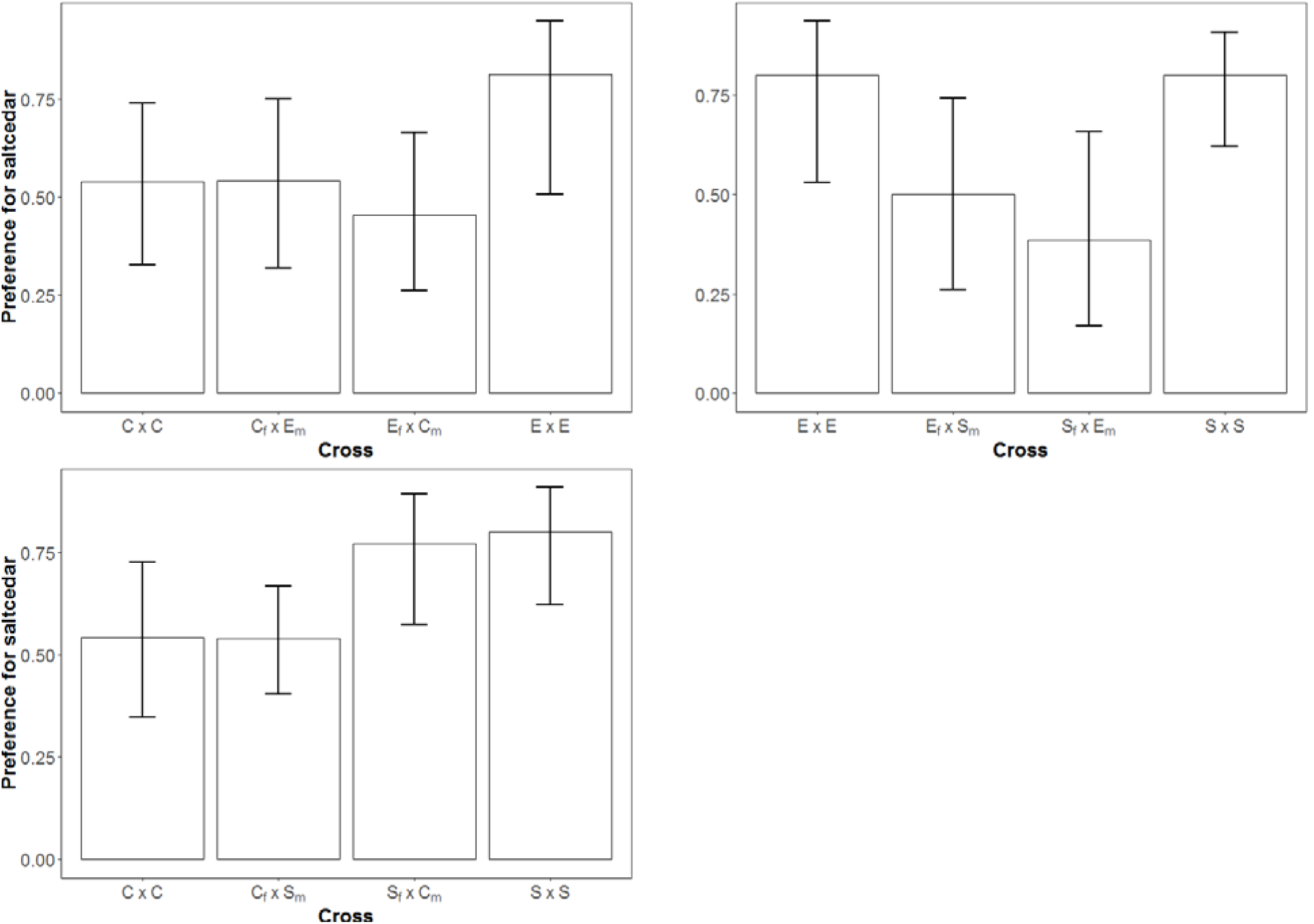
Preference for saltcedar (target) over athel (non-target) across all parental species (*D. carinata* “C”, *D. sublineata* “S”, *D. elongata* “E”) and their hybrids in the second generation of hybridization. Hybridization significantly affected host preference for the S × C and S × E crosses. Error bars represent 95% confidence intervals.

## 4. Discussion

In introduced species, the effects of hybridization can influence local adaption and determine the fate of colonization success and establishment (Rius & Darling 2014). Introduced biological control agents undergo similar pressures as newly invading species, and understanding the mechanisms behind population growth and establishment are crucial to the implementation of successful biological control. In this study, we investigated the effects of hybridization on various life history traits and host preference for three different species of the biological control agent *Diorhabda*. We confirmed that all three species were reproductively compatible (Bean et al. 2013b), and that found that reciprocal crosses produced viable offspring through at least two generations. Life history traits beyond the production of viable eggs were either unchanged or improved with hybridization when compared to the parental species. These results support the hypothesis that these species have not experienced reproductive isolation for long enough to allow the evolution of genetic incompatibilities.

Hybridization can have positive, neutral, or negative effects on fitness. These effects depend on the genetic distance between mixing populations and the interactions between genes and environment. Hybrid vigor is commonly seen in the first generation of admixture between genetically distinct populations, and is typically thought to be due to masking of deleterious alleles rather than overdominance (Szulkin *et al.* 2010), whereas hybrid breakdown is commonly seen in the second or later generations due to recombination of the parental genes, allowing for the possibility of deleterious allele combinations (heterozygote disadvantage) (Dobzhansky 1950; Edmands 2002). In our study, there was no difference between parents and their hybrid offspring in fecundity or percent hatching in the first generation in any cross. Previous molecular work done by Bean et al (2013b) showed that while all four *Diorhabda* species separated into their own clades, the three species examined here were likely more closely related to each other than to the congeneric *D. carinulata*. It is possible that these species are not genetically distinct enough to be detrimentally affected by hybridization. However, the beetles used in our study had been lab reared for varying amounts of time (at least ten generations), and may have become inbred or lost variation via drift, both of which could reduce fitness. Thus, an alternative explanation is that positive effects of crossing, via masking of deleterious mutations could have balanced out potentially negative effects of hybridization, leading to zero, or close to zero, net change in life history traits. The masking of deleterious mutations can persist for many generations (Frankham 2016; Hedrick & Garcia-Dorado 2016), and thus further study investigating the effects of hybridization for more than three generations would increase our understanding.

Our results show that some of our crosses benefited greatly from hybridization in fecundity and development time in the second generation, and thus we see no evidence of hybrid breakdown. S × C crosses produced 67% more eggs and developed approximately 7 days shorter than the parental species. The E × C cross exhibit the same trend, although this was only marginally significant. Other crosses showed no effect of hybridization, and none of our crosses suffered a fitness cost. In the S × C cross, where we could examine a third generation, we saw no effect of hybridization on fecundity, but we did see a trend that hybrids were developing slower on both host plants. For this analysis, our sample size was lower than for the previous generations, and so further work is necessary to determine if development time slowed because of hybridization.

Changes in host specificity in a released agent are one of the most concerning issues to scientists studying biological control (Van Klinken & Edwards 2002; Brodeur 2012; McEvoy *et al.* 2012). Our results show that host specificity can indeed be affected by hybridization, and that the phenotype can vary depending on the maternal or paternal species. In the S × C crosses, host preference of the hybrid followed the preference of the maternal species, whereas in the S × E cross, hybrids showed no preference for either host plant where the parents both showed a strong preference for the target host. Host specificity depends upon a suite of traits, such as behavior, morphology, and life-history strategies and as such is highly constrained (Zwolfer & Harris 1971; Giebink *et al.* 1984; Chang *et al.* 1987). Even so, in more generalist species than are typically used for biological control, host use has been shown to have a genetic basis, and can thus vary between individuals and populations (Singer & Parmesan 1993; Funk 1998). In our study, the inherited pattern for host use depended not only on the cross, but the preference of the maternal species. A growing body of literature suggests that for herbivorous insect species, mothers have been shown to influence host use (Amarillo-Suarez & Fox 2006; Egan *et al.* 2011; Cahenzli & Erhardt 2013). Egan *et al.* (2011) specifically demonstrated that host-use and performance are traits with sex-linked maternal influence. Consequently, the pattern of host specificity in hybrid crosses can be hard to predict since it will depend not only on the amount of genetic variation across a suite of traits, but also parental influence.

Using hybridization in biological control presents unique challenges. On one hand, increased genetic variation, potentially from hybridization, can buffer introduced populations against adaptive challenges and thus increase the probability of establishment and effective control (Hopper *et al.* 1993). On the other, the genetic admixture of previously isolated populations might give rise to new phenotypes that are less desirable, such as a change in host specificity (Hoffmann *et al.* 2002; Mathenge *et al.* 2010). Our results demonstrate that while some crosses benefit from hybridization in terms of development time and fecundity, differences in host specificity due to hybridization is of concern.

**Table 5:**
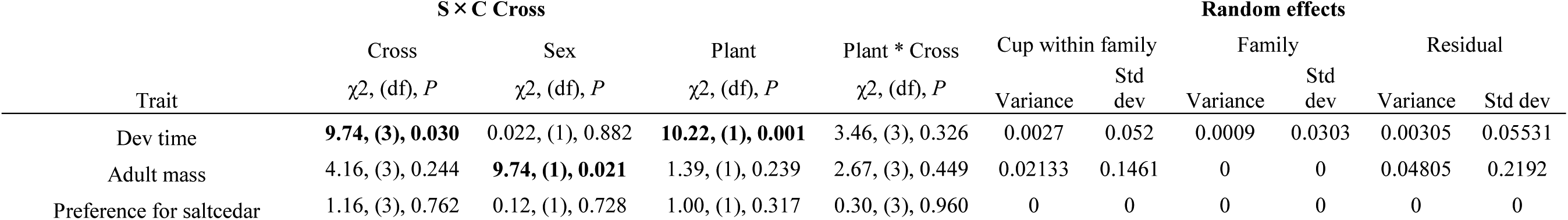
Results from generalized linear mixed-effects models for the third generation of the *D. sublineata* by *D. carinata* cross.

| Trait | **S × C Cross** |  |  |  |  | **Random effects** |  |  |  |  |
| --- | --- | --- | --- | --- | --- | --- | --- | --- | --- | --- |
|  | Cross | Sex | Plant | Plant * Cross | Cup within family |  | Family |  | Residual |  |
| | $\chi^2$, (df), *P* | $\chi^2$, (df), *P* | $\chi^2$, (df), *P* | $\chi^2$, (df), *P* | Variance | Std dev | Variance | Std dev | Variance | Std dev |
| Dev time | **9.74, (3), 0.030** | 0.022, (1), 0.882 | **10.22, (1), 0.001** | 3.46, (3), 0.326 | 0.0027 | 0.052 | 0.0009 | 0.0303 | 0.00305 | 0.05531 |
| Adult mass | 4.16, (3), 0.244 | **9.74, (1), 0.021** | 1.39, (1), 0.239 | 2.67, (3), 0.449 | 0.02133 | 0.1461 | 0 | 0 | 0.04805 | 0.2192 |
| Preference for saltcedar | 1.16, (3), 0.762 | 0.12, (1), 0.728 | 1.00, (1), 0.317 | 0.30, (3), 0.960 | 0 | 0 | 0 | 0 | 0 | 0 |

